# Effects of one night of sleep deprivation on single- and dual-task gait

**DOI:** 10.64898/2026.01.12.698966

**Authors:** Maura Seynaeve, Anna C Render, Peter C Fino, Toon de Beukelaar

## Abstract

Sleep deprivation impairs cognitive control, which may affect movements that rely on these processes, such as walking. To test whether gait changes after one night of sleep deprivation reflect reduced cognitive capacity, we compared its effects with those of dual-task walking (i.e., walking while performing a simultaneous cognitive task). We hypothesize that sleep deprivation will produce gait changes similar to those under dual-task conditions. Eighteen healthy adults (9 female, 9 male; 22.2 (2.3) yrs) were tested the morning after a sleep deprivation (SDEP) and a control night. Participants completed two 2-min trials: single-task walking and walking with a concurrent 2-back working memory task (DT, dual-task). Using lateral foot and pelvis marker trajectories, we calculated spatiotemporal parameters, foot placement error in antero-posterior (FPE_AP_) and mediolateral (FPE_ML_) directions, and mediolateral margin of stability (MoS_ML_). SDEP increased average step time (p<0.001) and step length (p=0.001), and DT reduced spatiotemporal variability. Both SDEP (p=0.001) and DT (p<0.001) reduced FPE_AP_, but only DT reduced FPE_ML_ (p<0.001). Additionally, mean MoS_ML_ decreased only in SDEP (p=0.011). Overall, these findings suggest that while sleep deprivation and dual-tasking both affect gait, the effects of sleep deprivation on gait cannot be fully explained by reduced cognitive resources.

## Introduction

Sleep is a biological process essential for maintaining physical health, cognitive function, and overall well-being across the lifespan. Besides the well-known effects of sleep loss on cognitive performance^1^, there is a growing body of literature showing that sleep loss also affects motor performance^2–6^. In general, these studies report a more profound effect of sleep loss for continuous and multi-joint tasks (e.g., maintaining balance) in comparison to discrete and single-joint tasks (e.g., a maximal knee extension). These continuous, multi-joint tasks rely on neural control processes that exceed basic spinal reflex mechanisms, including feedback and feedforward processes, executive control, and attention^7,8^. Such findings suggest that sleep loss disproportionately affects movements that rely heavily on these cognitive control processes. For instance, multiple studies have shown that sleep loss impairs balance^9,10^, a task that relies on these cognitive control processes^11,12^. However, most of these studies focus on static balance tasks, so it remains unclear whether sleep deprivation causes similar impairments during dynamic activities such as walking.

Observational studies support an association between sleep and gait. Individuals who report poor or insufficient sleep (<7 hours or poor sleep quality) show reduced gait speed, lower cadence and larger variability in spatiotemporal parameters^13^. Moreover, poor sleep is associated with increased trunk rotation, larger foot strike angles, longer stride length, and extended duration in single-leg support or swing phase, which collectively indicate compromised gait stability and efficiency^14–16^. However, the mechanisms underlying these relationships remain unclear because these prior studies have been purely observational and did not directly manipulate sleep.

Two interventional studies have experimentally altered sleep to examine the effects of sleep deprivation on gait, and these studies further support an association between the two. Oliveira et al. demonstrated that sleep deprivation reduces gait complexity, as reflected by a decrease in detrended fluctuation analysis (DFA) of inter-stride intervals. Thereby, the authors suggest sleep deprivation leads to altered neuromuscular control^17^. In addition, Umemura et al. reported that sleep-deprived participants exhibited impaired performance in a sensorimotor synchronization gait task, in which they were required to match their footfalls to auditory cues^18^. While these studies offer valuable insight into the sensorimotor mechanisms underlying gait, an alternative explanation for these relationships is a change in cognitive reserve after sleep deprivation.

Given the well-established finding that sleep deprivation severely depletes cognitive reserves^1^, it is possible that sleep-related alterations in gait stem from a decline in cognitive resources. One method to examine the contribution of these cognitive resources during gait is by introducing a dual-task interference paradigm^19^, where a cognitive task and a motor task are performed at the same time. Dual-tasking challenges the central nervous system’s ability to manage multiple demands and typically changes the task performance of one or both tasks. This interference can be explained by two distinct theoretical frameworks: bottleneck theory, which suggests that central processing for the second task is delayed until the first task is completed, or capacity-sharing theory, which proposes that central processing for both tasks can occur in parallel but must share limited cognitive resources. Similar to the association between poor sleep and gait, dual-tasking typically results in slower gait speeds and/or an increase in gait variability^20^. Therefore, the observed associations between gait disturbances and sleep might not be purely related to the motor system. Instead, gait alterations after insufficient sleep may represent the central nervous system’s difficulty in allocating sufficient cognitive resources to maintain normal gait.

To test this hypothesis, the present study aimed to examine the effects of one night of sleep deprivation on gait control and stability in single- and dual-task walking conditions. Utilizing a randomized, crossover design, participants underwent two testing sessions: one following a normal night of sleep (CON, control) and one following 24 hours of total sleep deprivation (SDEP). We hypothesized that sleep deprivation would produce effects similar to those observed under dual-task gait conditions, reflecting a shared underlying mechanism of reduced cognitive resource availability. In addition, we predict that the combination of sleep deprivation and dual-tasking would further exacerbate changes in gait parameters.

## Methodology

### Participants

Twenty-two healthy adults aged 18–35 years were recruited for the study. Sleep quality and daytime sleepiness were evaluated using the Pittsburgh Sleep Quality Index (PSQI) and the Epworth Sleepiness Scale (ESS), respectively. Only participants with a PSQI score ≤5 and an ESS score <11 were included to ensure adequate baseline sleep quality. Individuals with suspected or diagnosed sleep disorders were excluded. Additional exclusion criteria comprised daily alcohol consumption, caffeine intake exceeding 400 mg/day, use of psychoactive substances, active smoking, and any known metabolic, cardiovascular, respiratory, or orthopedic conditions that could influence task performance or sleep. Because a healthy sleep duration is defined as more than 7 hours, participants who slept less than 7 hours in the control condition were excluded. Based on this criterion, we had to exclude 4 participants, leaving 18 participants (9 female, 9 male; mean (sd) = 22.2 (2.3) yrs; 66.8 (9.1) kg; 1.74 (8.96) m) for data analysis. The study was approved by the Ethics Committee of UZ Leuven (S67790) and conducted in accordance with the Declaration of Helsinki. All participants provided written informed consent.

**Figure 1.**
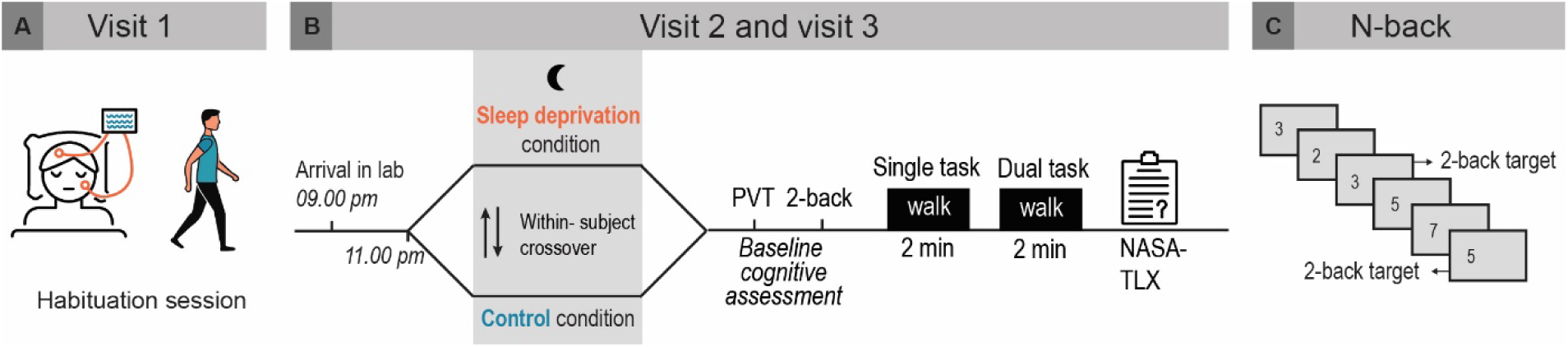
Study design and N-back task procedure. **A)** Participants first completed a habituation session (visit 1) to familiarize themselves with the sleep lab, polysomnography setup, and motor/cognitive tasks. **B)** In two subsequent experimental sessions (visits 2 and 3), participants either stayed awake overnight (sleep deprivation) or slept in the lab (control), in a randomized order. In the morning, baseline cognitive assessments included a psychomotor vigilance task (PVT) and a 2-back working memory task. Then, participants completed two 2-min treadmill trials: normal walking and walking while performing the 2-back task, followed by the NASA-TLX questionnaire to assess subjective task load. **C)** In the 2-back task, numbers 0–9 were presented, and participants responded when the current number matched the one presented two steps earlier.

### General study design

Seven days before the first experimental session, participants completed a habituation session during which they (i) slept in the laboratory to become accustomed to the sleep environment and (ii) were fully acquainted with the cognitive and motor tasks (fig. 1A). In the seven days before each of the two experimental sessions, participants were asked to maintain their normal sleep habits. During this period, physical activity and sleep were monitored using wrist-worn ActiGraph wGT3X-BT accelerometers (ActiGraph LLC, Pensacola, FL, USA) and the consensus sleep diary^21^. Participants were instructed to abstain from alcohol and caffeine before each experimental session.

Following the habituation session, participants completed two experimental sessions using a randomized, counterbalanced crossover design: one following a night of normal sleep (control condition, CON) and one following a night of total sleep deprivation (SDEP) (fig. 1B). Both conditions were interspersed by a one-week washout period. During the CON condition, participants slept in the laboratory from 11:00 p.m. to 7:00 a.m., with full polysomnographic (PSG) monitoring. In the SDEP condition, participants remained awake under continuous supervision throughout the night. They were seated in a dimly lit room (<30 lux) and allowed to engage in quiet activities such as reading or solving puzzles, with screen time limited to a maximum of 2 hours in total. Water was provided hourly, and small snacks (e.g., apple, yogurt, or biscuit) were offered at 11:00 p.m., 12:00 a.m., 1:00 a.m., and 2:00 a.m. The following morning, participants refrained from food intake but consumed a standardized 500 ml of water immediately upon waking (CON) or two hours prior to testing (SDEP). A series of cognitive and motor tasks commenced at 7:30 a.m. for CON and at 6:30 a.m. for SDEP.

### Cognitive and Motor tasks

#### Psychomotor vigilance task (PVT)

Sustained attention was assessed using a 10-minute Psychomotor Vigilance Task (PVT) performed at least 30 minutes after waking to minimize sleep inertia^22^. Participants responded as quickly as possible to visual stimuli presented at random intervals on a computer screen. Reaction times and lapses (responses >500 ms) were recorded.

#### N-back task

Working memory was evaluated using a 2-back task with numeric stimuli (fig. 1C)^23^. Participants were instructed to respond via button press whenever the current number matched the one presented two trials previously. Stimuli were presented at fixed 1.5-second intervals over a 2-minute testing period. Both response accuracy and reaction time were recorded.

#### Walking task

Participants then completed two 2-minute walking trials on an instrumented split-belt treadmill (Motekforce Link, Amsterdam, the Netherlands). Treadmill speed was set to each participant’s preferred walking speed as determined during the habituation session (mean (sd) = 1.25 (0.14) m/s). The first trial was performed without a cognitive task (single-task condition); participants were instructed to focus on a point ahead to minimize head movements and were asked to stay in the center of the treadmill. The second trial included the concurrent 2-back task (dual-task condition), with the numeric stimuli presented on a small screen mounted at eye level to the treadmill. Responses were recorded via a wired handheld button.

### NASA Task Load Index (NASA-TLX)

Upon completion of the testing protocol, participants’ perceived workload was assessed using the NASA-TLX questionnaire, which includes the mental demand, physical demand, frustration, performance, and effort subscales^24^.

### Instrumentation and data acquisition

#### Polysomnography and actigraphy

PSG data were collected using a mobile EEG system (LiveAmp, Brain Products, Gilching, Germany) at a sampling rate of 1000 Hz. EEG electrodes were placed at Fz, Cz, Pz, Oz, C3, and C4 according to the international 10–20 system, with the reference on the right mastoid and a ground electrode on the forehead. Vertical and horizontal eye movements were recorded using electrooculography (EOG) with electrodes above and below the right eye and at both outer canthi. Submental muscle activity was recorded via a chin electromyogram (EMG). EEG and EOG signals were band-pass filtered between 0.1 and 30 Hz, EMG signals between 10 and 200 Hz, and a 50 Hz notch filter was applied to all channels to reduce electrical interference.

Sleep staging was performed using the YASA toolbox in Python^25^, incorporating EEG (Cz), EOG, and EMG signals to classify wake, N1, N2, N3, and REM stages. Key sleep metrics including total sleep time, sleep efficiency, sleep onset latency, wake after sleep onset (WASO), number of awakenings, and stage durations were extracted for analysis.

Seven-day accelerometer data preceding the two experimental sessions were processed using the GGIR package in R (version 3.2-6)^26^. Raw acceleration signals were auto-calibrated, non-wear periods detected, and metrics of sleep and physical activity derived.

#### Marker data collection and preprocessing

Twenty-eight reflective markers were placed on anatomical bony landmarks and 12 technical cluster markers were placed on each thigh and shank. Thirteen infrared cameras (Vicon, Oxford Metrics, Oxford, UK) captured the positions of all reflective markers at a sampling rate of 250 Hz. We used OpenSim 4.5 (OpenSim, Stanford, USA) to first scale the Hamner musculoskeletal model based on the subject’s dimensions using a neutral standing trial^27^ and to subsequently export marker trajectories of the pelvis and lateral foot markers. The body’s center of mass (CoM) position was estimated as the average of the four pelvis markers located on the left and right anterior and posterior superior iliac spines^28^. All marker trajectories were low-pass filtered using a zero-phase 2nd-order Butterworth filter with a 6 Hz cutoff frequency. Filtered positions were then differentiated to compute marker velocity.

#### Gait event detection

Heel strikes (HS) and toe-offs (TO) were identified using the anteroposterior (AP) velocity of the lateral foot markers placed on the fifth metatarsophalangeal joints. HS events were defined as the transition from positive to negative AP velocity, and TO events as the transition from negative to positive velocity^29^. Stance phases were defined as periods between each HS and the subsequent TO. Midstance was defined at 50 percent of the step. Across all participants, an average (sd) of 222.0 (15.72) steps per trial were analyzed.

### Biomechanical outcome measures

#### Spatiotemporal parameters

We calculated step time, step length, and step width for each step to characterize participants’ general gait patterns. Step time was defined as the temporal interval between successive heel strikes of alternating feet. Step length was calculated using the AP positions of the leading and stance feet at contralateral toe-off. Similarly, step width was calculated based on the mediolateral distance between lateral foot markers on the leading and stance foot at contralateral toe-off. We calculated the within-trial mean and standard deviation of step time, step length, and step width from their respective time series.

#### Foot placement control

Based on Wang and Srinivasan, a multiple linear regression model was used to assess lateral foot placement control (FP_ML_)^30^. The mediolateral position (CoM_pos_ML_) and velocity (CoM_vel_ML_) of the CoM at midstance were included as independent variables, while the ML position of the subsequent contralateral foot (FP_ML_) at heel strike was included as the dependent variable. Both predictors and outcomes were expressed relative to the ipsilateral stance foot and mean-centered prior to analysis. The degree of foot placement control was quantified as the relative explained variance (R^2^) of the linear regression. Foot placement error (FPE) was quantified as the root mean square of the regression residuals, providing a measure of deviation from the predicted foot placement and thus precision of foot placement.

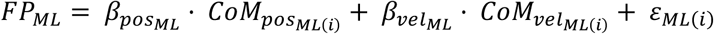

Similar to the model for lateral foot placement, a multiple linear regression model for the AP foot placement was fitted. In this case, AP position of the subsequent contralateral foot (FP_AP_) was predicted based on the AP position (CoM_pos_AP_) and velocity of the CoM (CoM_vel_AP_) at ipsilateral TO.

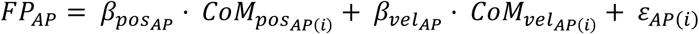

#### Mediolateral Margins of stability (MoS)

Dynamic margins of stability (MoS_ML_) were calculated based on the extrapolated center of mass (XcoM) concept^31^. The XcoM was defined as:

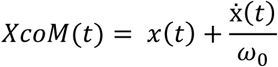

where *x* is the COM position, *x* is the COM velocity, and ω_₀_ is the natural frequency of the inverted pendulum:

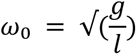

where 𝑔 = 9.81 𝑚/𝑠^2^ representing gravitational acceleration, and *l* the effective pendulum length, defined as the mean distance from the lateral ankle to the greater trochanter marker during the static trial.

The mediolateral MoS was then defined as the distance between the XcoM and the boundary of the base of support (BOS):

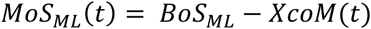

The boundary of BOS was defined by the mediolateral position of the lateral stance foot marker. Consistent with recommendations, the MoS was extracted at contralateral TO for each step^32^ and multiplied by a -1 for alternating steps such that a positive MoS_ML_ values indicated that the extrapolated center of mass remains within the base of support, reflecting mechanically stable steps, whereas negative values indicate that the center of mass extends beyond the base of support, reflecting instantaneous mechanical instability. Mean and variability in MoS_ML_ were quantified as the mean and standard deviation from the MOS_ML_ values across all steps within each trial.

### Statistics

Linear mixed-effects models evaluated the effects of sleep condition (CON vs. SDEP) and task (single- vs. dual-task) on all biomechanical outcome measures and N-back task performance. For the N-back analyses, trial type reflected whether the assessment was conducted in the morning (seated, no walking) or during the dual-task walking condition. Models included fixed effects for condition and trial, their two-way interaction, and random effects for subjects. The factor subjects were modeled with a random intercept to account for repeated measures within individuals. Pairwise comparisons between each level of the fixed effects were conducted, and Cohen’s d was calculated to estimate pair-wise effect sizes (see supplementary table 1). In addition, NASA-TLX and PVT scores were analyzed using paired-samples t-tests. All analyses were performed in R using the lme4 and lmerTest packages. Data are presented as mean (sd), and the significance level was set at p<0.05.

## Results

### Sleep architecture

PSG recordings during the control night indicated a mean total sleep duration of 7.66 (0.51) hours, with an average sleep efficiency of 84.24 (4.14) %. Wake after sleep onset (WASO) averaged 42.71 (19.45) minutes. The proportional distribution of sleep stages was: 6.27 (2.98) % for non-rapid eye movement (NREM) stage 1 sleep, 51.19 (5.54) % for stage 2, 19.41 (3.71) % for stage 3, and 23.14 (3.89) % of total sleep time for rapid eye movement (REM) sleep. These values are consistent with normative ranges, indicative of high-quality nocturnal sleep ^33^.

**Figure 2.**
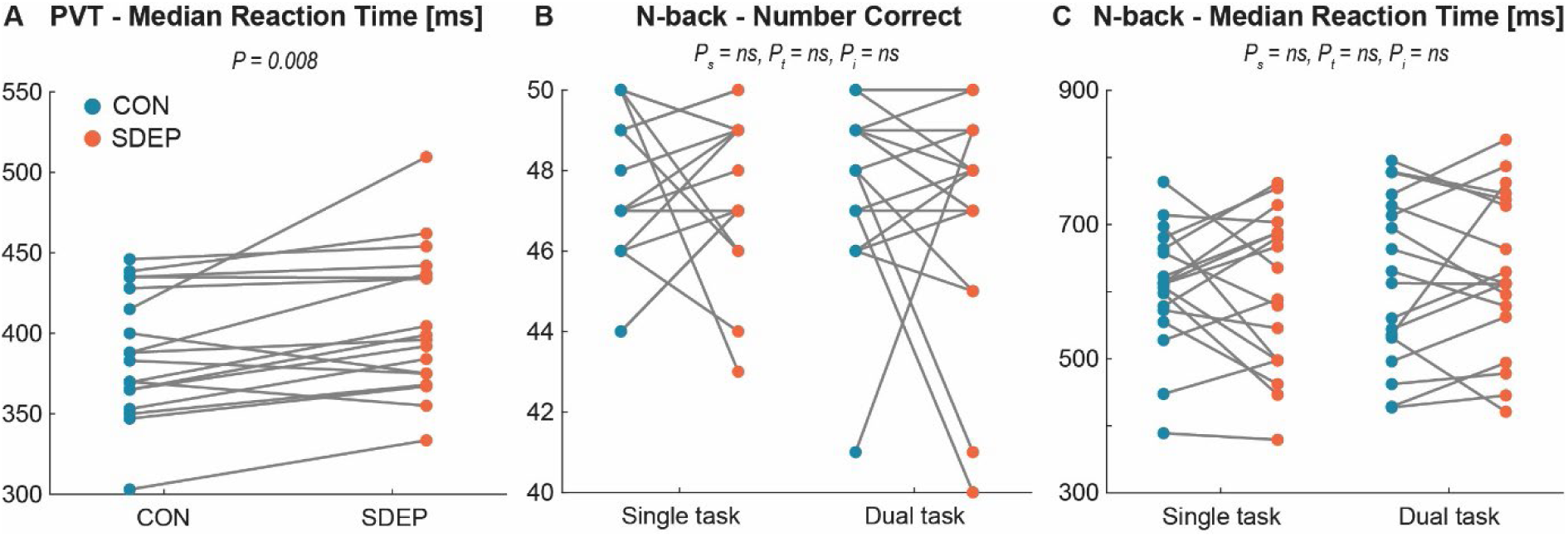
Performance on the cognitive tasks for the control condition (CON) in blue and sleep deprivation condition (SDEP) in red. A) Median reaction time on the PVT task. B) Number of correct responses (maximal score = 50) and, C) median reaction time during the N-back task in single-task condition (seated, in the morning) and dual-task condition (during walking). Each dot represents a participant, with lines linking repeated measurements from the same individual. P_s_ = main effect of sleep condition, P_t_ = main effect of task, P_i_ = sleep condition x task interaction effect.

### Cognitive tasks

Median reaction time on the PVT task (fig. 2A) was longer following SDEP compared with CON (SDEP: 407 (44.4) ms, CON: 388 (39.2) ms, *t*(17)=–3.02, p=.008, Cohen’s d=0.711). The N-back task was administered both in the morning as a single-task and during walking as a dual-task. No differences were observed in the number of correct responses (fig. 2B; maximal score=50) or in median reaction times (fig. 2C) between the sleep conditions or between the single- and dual-task assessment.

### Spatiotemporal parameters

**Figure 3.**
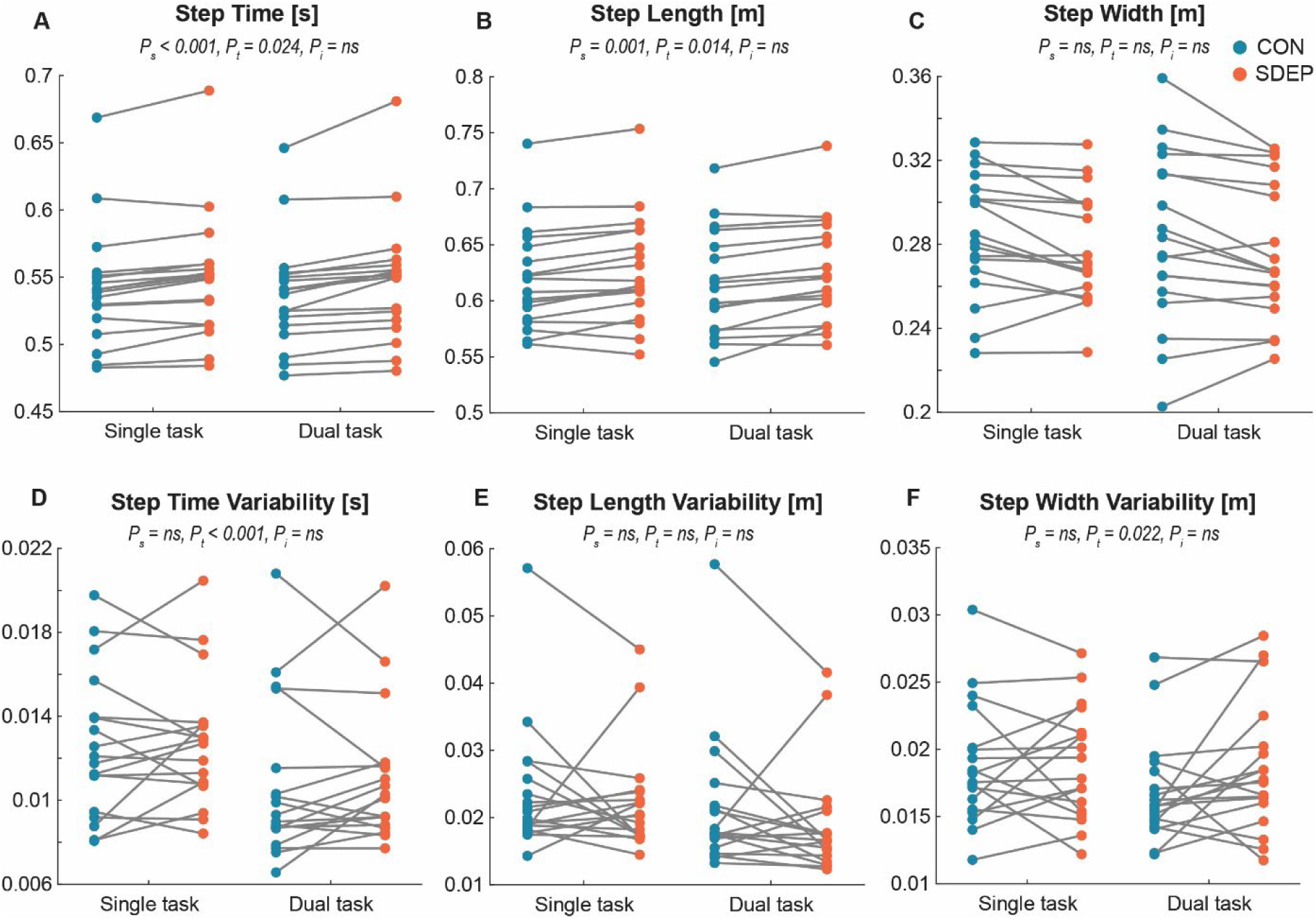
Spatiotemporal walking parameters under control (CON; blue) and sleep deprivation (SDEP; red) conditions. Data are shown for single-task walking (no cognitive task) and dual-task walking (simultaneous N-back performance). Each dot represents an individual participant, with lines connecting repeated measurements from the same participant. P_s_ = main effect of sleep condition, P_t_ = main effect of task, P_i_ = sleep condition x task interaction effect.

SDEP and DT both influenced spatiotemporal gait parameters, but in distinct ways. SDEP led to a small but significant increase in mean step time (p<0.001, fig. 3A) and step length (p=0.001, fig. 3B), while step time and step length variability (fig. 3D and fig. 3E), as well as step width and its variability (fig. 3C and fig. 3F), were unaffected. In contrast, performing the N-back task while walking resulted in shorter (p=0.014) and faster steps (p=0.024), along with reduced step time variability (p<0.001) and reduced step width variability (p=0.022), suggesting more consistent stepping under cognitive load. No significant interactions between sleep condition and task were observed for any of the spatiotemporal measures (all p>0.05). Results of the linear mixed-model and descriptive statistics can be found in Tables 1 and 2.

### Foot placement prediction

**Figure 4.**
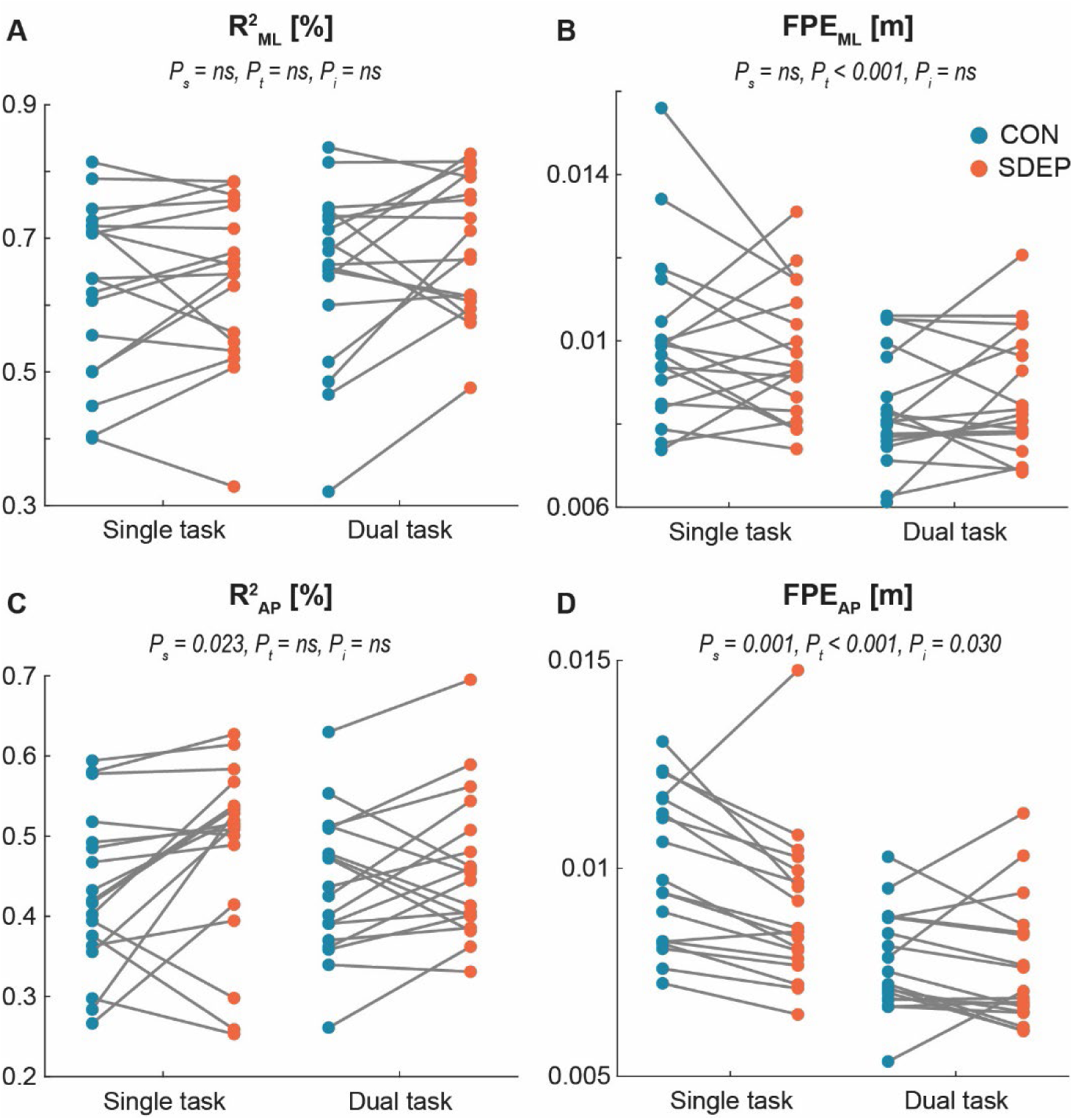
R^2^ and foot placement error (FPE) in mediolateral (ML) and anteroposterior (AP) directions under control (CON; blue) and sleep deprivation (SDEP; red) conditions. Data are shown for single-task walking (no cognitive task) and dual-task walking (simultaneous N-back performance). Each dot represents an individual participant, with lines connecting repeated measurements from the same participant. P_s_ = main effect of sleep condition, P_t_ = main effect of task, P_i_ = sleep condition x task interaction effect.

SDEP and DT uniquely affected ML and AP foot placement control. For R²_ML_, neither SDEP (p=0.528) nor DT (p=0.226) had a significant effect, and the interaction was non-significant (p=0.338, fig. 4A). However, FPE_ML_ was smaller during DT compared to single-task walking (p<0.001), but SDEP had no effect (p=0.363), and the interaction remained non-significant (p=0.166, fig. 4B).

In the AP direction, SDEP increased R² (p=0.023), whereas dual-tasking had no effect (p=0.699) and the interaction was non-significant (p=0.351, fig. 4C). FPE_AP_ was reduced by both SDEP (p=0.001) and DT (p<0.001). A small but significant interaction (p=0.030, fig. 4D) was present, showing that the effect of adding a dual-task had a larger effect on FPE_AP_ in the CON condition compared to the SDEP condition.

### Margins of stability

**Figure 5.**
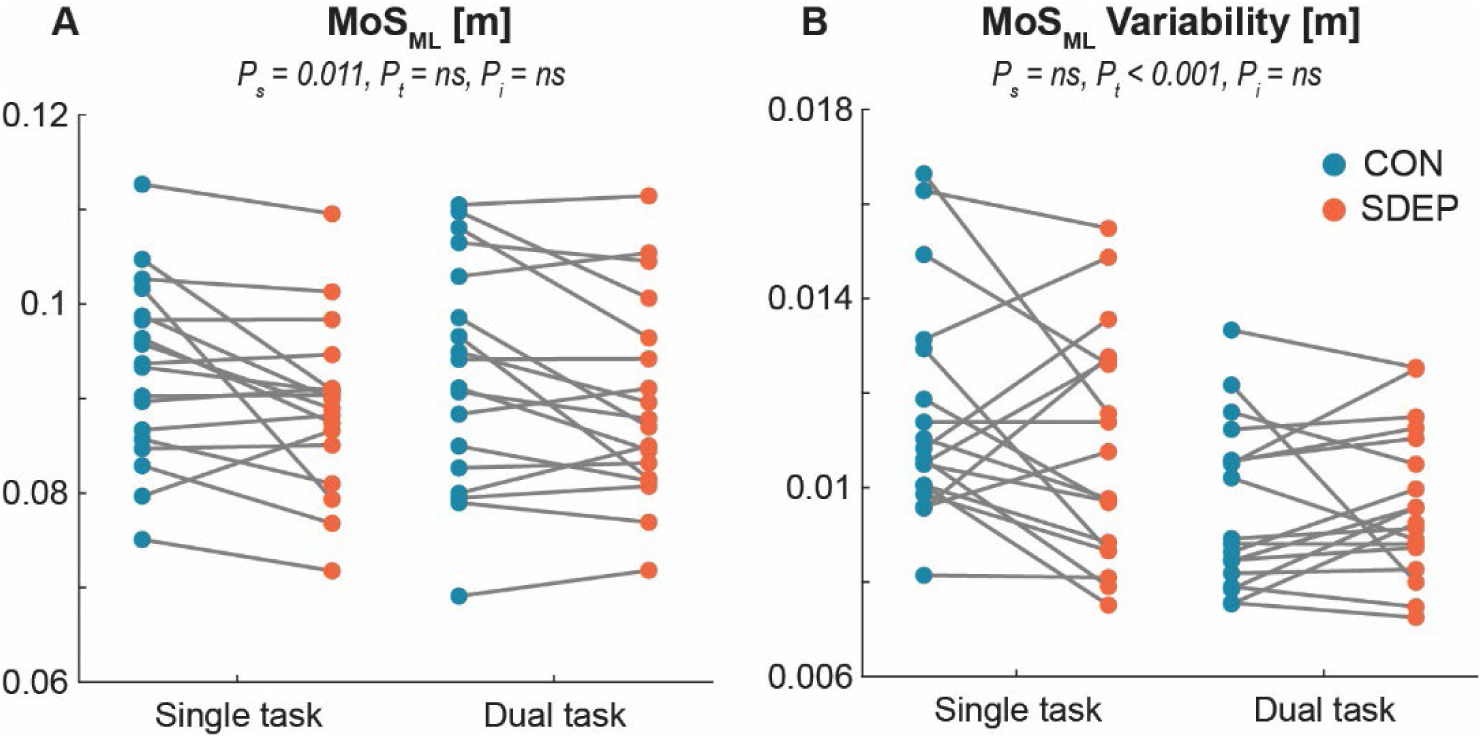
Mean and variability (SD) of margin of stability (MoS_ML_) under control (CON; blue) and sleep deprivation (SDEP; red) conditions. Data are shown for single-task walking (no cognitive task) and dual-task walking (simultaneous N-back performance). Each dot represents an individual participant, with lines connecting repeated measurements from the same participant. P_s_ = main effect of sleep condition, P_t_ = main effect of task, P_i_ = sleep condition x task interaction effect.

SDEP led to a decrease in mean MoS_ML_ (p=0.011), indicating slightly reduced lateral stability, whereas DT did not significantly affect mean MoS_ML_ (p=0.815, fig. 5A). There was no significant interaction between sleep condition and task (p=0.650). In terms of MoS_ML_ variability, DT reduced variability (p<0.001), while SDEP had no significant effect (p=0.074), and the interaction remained non-significant (p=0.157, fig. 5B). These results suggest that SDEP subtly decreases stability, whereas cognitive load reduced variability without changing the average MoS_ML_.

**Table 1.**
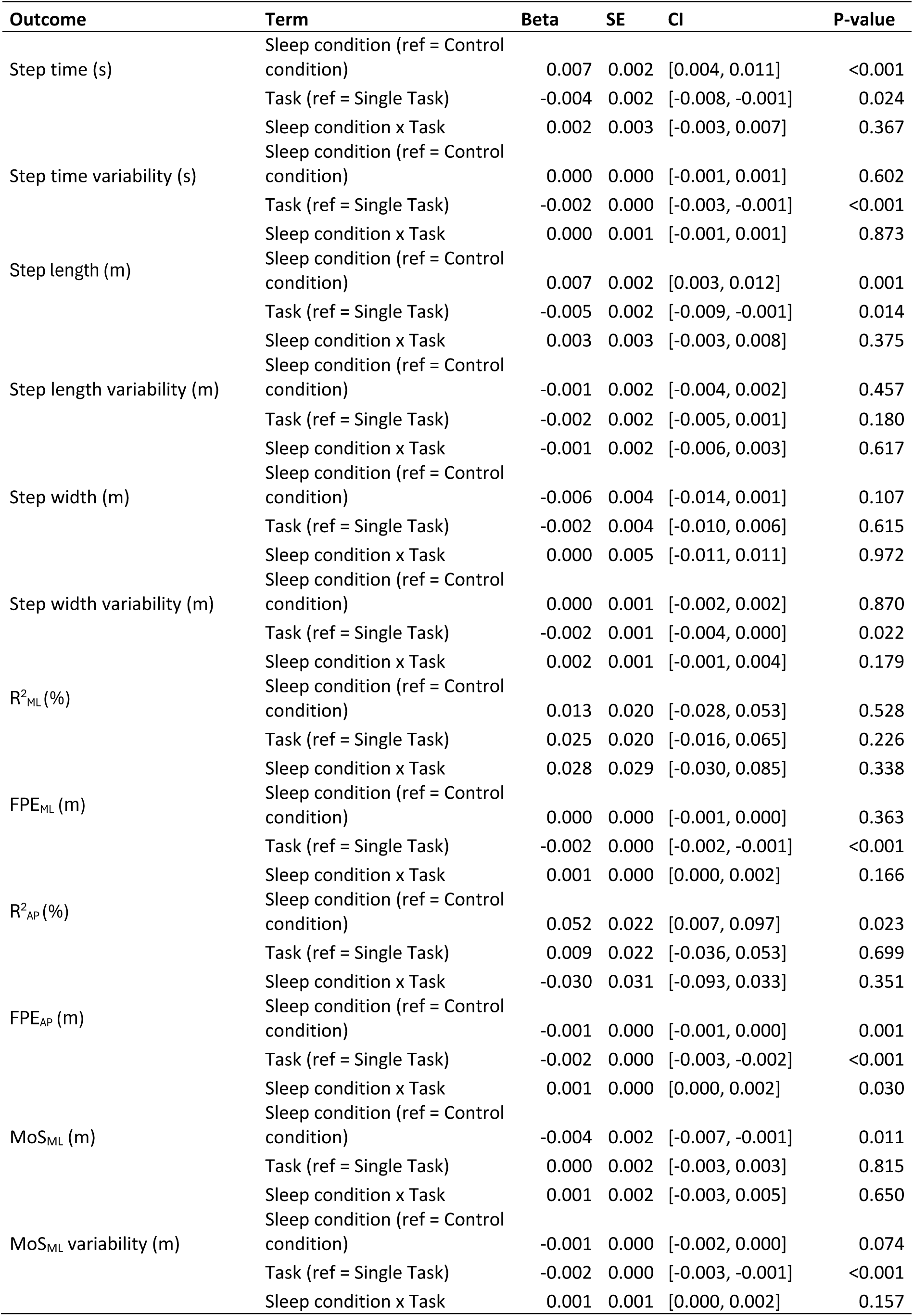
Linear mixed-model results for each outcome, including beta coefficients, standard errors, 95% confidence intervals (CIs), and p-values.

| Outcome | Term | Beta | SE | CI | P-value |
| --- | --- | --- | --- | --- | --- |
| Step time (s) | Sleep condition (ref = Control condition) | 0.007 | 0.002 | [0.004, 0.011] | <0.001 |
|  | Task (ref = Single Task) | -0.004 | 0.002 | [-0.008, -0.001] | 0.024 |
|  | Sleep condition x Task | 0.002 | 0.003 | [-0.003, 0.007] | 0.367 |
| Step time variability (s) | Sleep condition (ref = Control condition) | 0.000 | 0.000 | [-0.001, 0.001] | 0.602 |
|  | Task (ref = Single Task) | -0.002 | 0.000 | [-0.003, -0.001] | <0.001 |
|  | Sleep condition x Task | 0.000 | 0.001 | [-0.001, 0.001] | 0.873 |
| Step length (m) | Sleep condition (ref = Control condition) | 0.007 | 0.002 | [0.003, 0.012] | 0.001 |
|  | Task (ref = Single Task) | -0.005 | 0.002 | [-0.009, -0.001] | 0.014 |
|  | Sleep condition x Task | 0.003 | 0.003 | [-0.003, 0.008] | 0.375 |
| Step length variability (m) | Sleep condition (ref = Control condition) | -0.001 | 0.002 | [-0.004, 0.002] | 0.457 |
|  | Task (ref = Single Task) | -0.002 | 0.002 | [-0.005, 0.001] | 0.180 |
|  | Sleep condition x Task | -0.001 | 0.002 | [-0.006, 0.003] | 0.617 |
| Step width (m) | Sleep condition (ref = Control condition) | -0.006 | 0.004 | [-0.014, 0.001] | 0.107 |
|  | Task (ref = Single Task) | -0.002 | 0.004 | [-0.010, 0.006] | 0.615 |
|  | Sleep condition x Task | 0.000 | 0.005 | [-0.011, 0.011] | 0.972 |
| Step width variability (m) | Sleep condition (ref = Control condition) | 0.000 | 0.001 | [-0.002, 0.002] | 0.870 |
|  | Task (ref = Single Task) | -0.002 | 0.001 | [-0.004, 0.000] | 0.022 |
|  | Sleep condition x Task | 0.002 | 0.001 | [-0.001, 0.004] | 0.179 |
| $R^2_{ML}$ (%) | Sleep condition (ref = Control condition) | 0.013 | 0.020 | [-0.028, 0.053] | 0.528 |
|  | Task (ref = Single Task) | 0.025 | 0.020 | [-0.016, 0.065] | 0.226 |
|  | Sleep condition x Task | 0.028 | 0.029 | [-0.030, 0.085] | 0.338 |
| $FPE_{ML}$ (m) | Sleep condition (ref = Control condition) | 0.000 | 0.000 | [-0.001, 0.000] | 0.363 |
|  | Task (ref = Single Task) | -0.002 | 0.000 | [-0.002, -0.001] | <0.001 |
|  | Sleep condition x Task | 0.001 | 0.000 | [0.000, 0.002] | 0.166 |
| $R^2_{AP}$ (%) | Sleep condition (ref = Control condition) | 0.052 | 0.022 | [0.007, 0.097] | 0.023 |
|  | Task (ref = Single Task) | 0.009 | 0.022 | [-0.036, 0.053] | 0.699 |
|  | Sleep condition x Task | -0.030 | 0.031 | [-0.093, 0.033] | 0.351 |
| $FPE_{AP}$ (m) | Sleep condition (ref = Control condition) | -0.001 | 0.000 | [-0.001, 0.000] | 0.001 |
|  | Task (ref = Single Task) | -0.002 | 0.000 | [-0.003, -0.002] | <0.001 |
|  | Sleep condition x Task | 0.001 | 0.000 | [0.000, 0.002] | 0.030 |
| $MoS_{ML}$ (m) | Sleep condition (ref = Control condition) | -0.004 | 0.002 | [-0.007, -0.001] | 0.011 |
|  | Task (ref = Single Task) | 0.000 | 0.002 | [-0.003, 0.003] | 0.815 |
|  | Sleep condition x Task | 0.001 | 0.002 | [-0.003, 0.005] | 0.650 |
| $MoS_{ML}$ variability (m) | Sleep condition (ref = Control condition) | -0.001 | 0.000 | [-0.002, 0.000] | 0.074 |
|  | Task (ref = Single Task) | -0.002 | 0.000 | [-0.003, -0.001] | <0.001 |
|  | Sleep condition x Task | 0.001 | 0.001 | [0.000, 0.002] | 0.157 |

**Table 2.** Descriptive statistics (mean (SD)) for each outcome by task and sleep condition.

| Outcome | Single-task |  | Dual-task |  |
| --- | --- | --- | --- | --- |
|  | CON | SDEP | CON | SDEP |
| Step time (s) | 0.542 (0.044) | 0.549 (0.046) | 0.538 (0.041) | 0.547 (0.046) |
| Step time variability (s) | 0.013 (0.003) | 0.013 (0.003) | 0.011 (0.004) | 0.011 (0.003) |
| Step length (m) | 0.620 (0.046) | 0.627 (0.048) | 0.615 (0.046) | 0.625 (0.045) |
| Step length variability (m) | 0.024 (0.010) | 0.022 (0.008) | 0.021 (0.010) | 0.019 (0.008) |
| Step width (m) | 0.285 (0.029) | 0.278 (0.026) | 0.283 (0.041) | 0.276 (0.033) |
| Step width variability (m) | 0.019 (0.005) | 0.019 (0.004) | 0.017 (0.004) | 0.019 (0.005) |
| $R^2_{ML}$ (%) | 0.625 (0.130) | 0.637 (0.121) | 0.649 (0.130) | 0.690 (0.104) |
| $FPE_{ML}$ (m) | 0.010 (0.002) | 0.010 (0.002) | 0.008 (0.001) | 0.009 (0.001) |
| $R^2_{AP}$ (%) | 0.429 (0.100) | 0.481 (0.113) | 0.438 (0.089) | 0.460 (0.092) |
| $FPE_{AP}$ (m) | 0.010 (0.002) | 0.009 (0.002) | 0.008 (0.001) | 0.008 (0.002) |
| $MoS_{ML}$ (m) | 0.093 (0.093) | 0.089 (0.090) | 0.010 (0.012) | 0.009 (0.011) |
| $MoS_{ML}$ variability (m) | 0.012 (0.010) | 0.011 (0.010) | 0.002 (0.002) | 0.002 (0.002) |

### NASA-TLX

**Table 3.** Results of paired samples t-test and descriptive statistics (mean (SD)) of the NASA-TLX by sleep condition.

| Outcome | CON |  | SDEP |  | diff | t | P-value | Cohen's d |
| --- | --- | --- | --- | --- | --- | --- | --- | --- |
| Mental Demand | 8.167 | (4.902) | 10.000 | (5.831) | 1.833 | -1.838 | 0.084 | 0.335 |
| Physical Demand | 1.889 | (1.676) | 3.500 | (4.260) | 1.611 | -1.767 | 0.095 | 0.449 |
| Performance | 3.556 | (2.975) | 4.111 | (2.867) | 0.556 | -0.627 | 0.539 | 0.190 |
| Effort | 7.778 | (4.977) | 11.500 | (4.878) | 3.722 | -3.908 | 0.001 | 0.755 |
| Frustration | 4.389 | (4.539) | 7.333 | (5.347) | 2.944 | -2.773 | 0.013 | 0.588 |
| Total NASA-TLX | 25.778 | (13.371) | 36.444 | (15.912) | 10.667 | -4.035 | 0.001 | 0.712 |

*Diff, absolute difference between values of control condition (CON) and sleep deprivation condition (SDEP).*

Subjective workload was assessed using the NASA-TLX immediately after the dual-task walking condition (Table 3). Overall workload, as indicated by the Total NASA-TLX score, was higher during SDEP compared with the control condition (p<0.001). Among the subscales, participants reported greater effort (p=0.001) and frustration (p=0.013) during SDEP. Mental demand (p=0.083) and physical demand (p=0.095) showed trends toward higher ratings in SDEP, although these did not reach statistical significance. Performance ratings did not differ significantly between conditions (p=0.539).

## Discussion

The present study investigated the effects of one night of total sleep deprivation on gait control and stability under single- and dual-task walking conditions. We hypothesized that sleep deprivation would produce effects similar to dual-task walking, reflecting a shared underlying mechanism of reduced cognitive resource availability. In addition, we expected the impact of sleep deprivation to be larger in the dual-tasking condition as observed by a significant interaction effect. However, comparing the effects of sleep deprivation and dual-tasking showed mixed results. To obtain a comprehensive view of gait adaptations, we examined gait using three complementary analytical approaches. At the spatiotemporal level, sleep deprivation increased step time, whereas dual-tasking decreased it. In addition, dual-tasking primarily reduced gait variability, whereas sleep deprivation did not affect variability. These measures describe overall changes in gait behavior but do not directly reflect underlying control strategies. Therefore, we next examined foot placement control, given its key role in stabilizing gait^30^. We found decreased FPE_AP_ under both dual-tasking and sleep deprivation, whereas FPE_ML_ was only affected by dual-tasking. Finally, to determine whether these changes translated into altered balance control, we assessed mediolateral margins of stability. MoS_ML_ was reduced following sleep deprivation but remained unaltered during dual-tasking, suggesting that despite changes in spatiotemporal parameters and foot placement control in both conditions, only sleep deprivation resulted in measurable impairments in mediolateral instantaneous stability. Overall, the present results suggest the effects of dual-tasking and sleep deprivation on gait arise from distinct underlying mechanisms. More specifically, sleep deprivation might (i) impact different subcomponents of cognitive functions compared to dual-tasking and/or (ii) affect gait through mechanisms that extend beyond cognitive factors.

### Subcomponents of attention

Although both sleep deprivation and dual-tasking impact attention, they might do so by affecting different subcomponents of attention. The differential effects of sleep deprivation and dual-tasking can be understood using van Zomeren and Brouwer’s multicomponent model of attention, which distinguishes intensity and selectivity dimensions^34^. Sleep deprivation primarily compromises the *intensity* components, i.e. tonic alertness and sustained vigilance, reducing overall arousal and producing a global reduction in the system’s capacity. This effect manifests as pronounced impairments in basic, continuous processing tasks, while higher-order functions such as working memory and reasoning are relatively spared^35–38^. Consistent with this, participants in the present study showed increased median reaction times on the PVT following total sleep deprivation (fig. 2A), whereas performance on the 2-back task remained stable (fig. 2B). In contrast, dual-tasking challenges the *selectivity* dimension by requiring the flexible allocation and coordination of attentional resources across concurrent activities via executive control. Performance decrements under dual-tasking therefore arise from competition for shared cognitive resources within an otherwise alert system. Taken together, these findings indicate that sleep deprivation reduces baseline alertness and attentional capacity (intensity deficit), whereas dual-tasking interferes with the distribution of attention among competing tasks (selectivity deficit).

### Gait adaptations under dual-tasking

Given that attention and executive functions are necessary for stable gait, impairments in both types of attention may pose a threat to stability and negatively impact gait^39^. When dynamic stability during gait is challenged (e.g., by perturbations or uneven terrain), individuals typically adopt a protective gait pattern characterized by shorter and wider steps^40,41^. Increasing step width enhances the lateral margin of stability, while shorter steps help maintain the center of mass within this margin and reduce vulnerability to slips or trips^42^. Thus, shorter and wider steps are generally interpreted as a stability-enhancing strategy. The strategy of increasing step width has also been reported in clinical populations, which has been interpreted as a compensatory response to counteract the impaired postural control associated with these disorders^43,44^.

This protective pattern was somehow evident in the dual-task condition in the current study. Apart from taking shorter steps (fig. 3A), sleep-deprived participants also reduced their spatiotemporal variability (fig. 3D and 3F). Gait variability reflects the extent to which the motor system explores different movement solutions across steps. Reduced spatiotemporal variability indicates tighter neuromotor regulation, often described as a stiffer control strategy, which limits adaptability but reduces unexpected deviations that could compromise balance. This stiffer control strategy is also reflected in the lower FPE in both the AP and ML directions (fig. 4B and 4D). FPE quantifies how closely actual foot placement matches the placement predicted to regulate the center of mass. Lower FPE indicates tighter coupling between foot placement and the instantaneous CoM state, reflecting reduced tolerance for deviation and increased reliance on predictive, tightly regulated control rather than adaptive or exploratory behavior. Taken together, these adaptations could be interpreted as a stiffer, less adaptable, yet seemingly “safer” walking pattern.

Alternatively, the gait adaptation during dual-tasking may reflect a shift toward a more automatic control strategy in response to the reduced attentional selectivity. Because dual-tasking taxes executive functions responsible for task coordination and flexible resource allocation, actively controlling gait becomes more costly. To mitigate this executive load, the motor system may adopt a simplified and more stereotyped gait pattern that relies more heavily on automatic, subcortical control processes and less on continuous executive supervision. Doing so, cognitive resources can be reallocated toward the secondary cognitive task. This interpretation is supported by postural control studies in young adults showing reduced sway while standing when a dual-task was added^45–48^. Importantly, these reductions occurred without increases in ankle muscle activity, indicating that stabilization is not achieved through stiffening but rather through greater reliance on automatic control processes^49^. These studies therefore suggest that the shift from an internal to an external focus during dual-tasking can enhance the automaticity and stability of postural control. Thus, in this population, dual-tasking may not necessarily pose a threat to gait stability or lead to a protective gait pattern. Instead, dual-tasking may be associated with a shift toward a more automatic gait pattern that helps preserve stability.

### Gait adaptations under sleep deprivation

In contrast to dual-tasking, participants in the sleep deprivation condition showed reduced instantaneous gait stability as observed by their decreased average lateral MoS (fig. 5A). Based on their increased step length and step time (fig. 3A and 3B), they were not adopting the protective gait pattern as described above. Rather, these adaptations may reflect an effort to conserve energy and walk as passively as possible. Walking with shorter and wider steps is metabolically costly, indicating that optimizing stability carries an energetic penalty^50–52^. In line with this, the observed decrease in FPE_AP_ (fig. 4D) following sleep deprivation could reflect a strategy to limit push-off torque around the ankle joint, as foot placement errors in the AP direction are compensated by increasing push-off torque in the ankle^53^. Therefore, sleep-deprived participants might try to limit their errors in an attempt to avoid the increased torques around the ankle joint. In addition, the increased R^2^_AP_ (fig. 4C) indicates that participants relied more heavily on their CoM state during foot placement, signifying a shift toward more passive gait dynamics^30^. Together, these gait adaptations came at the cost of instantaneous gait stability.

Despite the reduced instantaneous lateral stability, participants did not exhibit active adjustments in FPE_ML_ or step width (fig. 4B and 3C) that might have helped preserve stability. Sleep deprivation is known to reduce alertness, which may compromise feedback control by delaying or weakening the detection and correction of gait errors. Reduced alertness may also impair feedforward mechanisms by limiting the ability to update and adapt predictive motor plans based on recent sensory information. This interpretation is in line with research showing that feedback and feedforward processes of motor control are indeed impacted by sleep loss. In the study of Umemura et al., participants walked on a treadmill while synchronizing their footsteps with auditory cues after one night of sleep deprivation^18^. They observed a significant increase in period errors, i.e. the difference in duration between two consecutive steps and two consecutive auditory cues, which the authors link to deficits in feedforward control^54^. In another study, decreased performance on a visually guided force-control task was found after sleep restriction^55^. As this task depends on sustained attention and feedback-based corrections, its impairment provides additional evidence that feedback-related sensorimotor processes are disrupted by sleep loss. Thus, the attentional intensity deficit induced by sleep deprivation may impair the ability to generate active feedback and feedforward-based corrections, especially in the ML plane. Effective corrections in the ML plane require a heightened level of control and vigilance compared to the AP plane, which may explain why FPE was adjusted in the AP plane but remained unchanged in the ML plane^56–58^. This lack of active corrections and passive gait dynamics under sleep deprivation might make subjects more susceptible to perturbations. Future studies should incorporate external perturbations to directly test this hypothesis.

### Limitations

Several limitations should be considered when interpreting the results of this study. First, although each trial included a sufficient number of steps for reliable gait analysis, the relatively short trial duration may have limited the expression of sleep deprivation–related gait alterations. Participants may have been able to temporarily compensate and maintain performance over the brief testing period, whereas longer or repeated trials might reveal effects associated with accelerated mental or physical fatigue induced by sleep deprivation. Second, the absence of changes in cognitive task performance suggests that the choice of cognitive task may not have been optimal for detecting the effects of sleep deprivation. Sleep deprivation has been shown to disproportionately impair sustained attention compared with certain aspects of executive control^35–38^, and the task used in this study may not have sufficiently taxed this sustained attention. Alternatively, some studies suggest that individuals undergoing sleep deprivation may allocate more compensatory effort to easier tasks and choose to detach from more challenging tasks^59^. Therefore, the task difficulty in this study may have been too low, allowing participants to maintain performance through compensatory strategies. Third, the fixed order of the single-task and dual-task conditions represents an important limitation when comparing performance across tasks. Because task order was not counterbalanced, potential order effects, such as treadmill habituation or fatigue, cannot be disentangled from true task-related differences. Fourth, it is important to note that correlations between the center of mass (CoM) state and foot placement alone cannot determine whether the observed relationship reflects active control mechanisms or passive biomechanical dynamics^60^. While such correlations provide insight into gait patterns, they do not by themselves reveal whether adjustments are the result of deliberate, feedback-driven control or emergent passive stability strategies. Fifth, participants slept in the laboratory during the control condition, which could have reduced sleep quality compared to a typical night at home, potentially limiting the validity of comparisons between conditions. Finally, while acute sleep deprivation provides a controlled experimental model, it is less representative of the chronic, partial sleep restriction commonly experienced in real-world settings, which may reduce the ecological validity of the findings.

## Conclusions

In summary, the present study demonstrates that one night of total sleep deprivation and dual-tasking have distinct effects on gait control and stability. Dual-tasking led to a reduction in step time, step length, spatiotemporal variability and foot placement error in both mediolateral as well as anteroposterior planes, while preserving instantaneous gait stability. In contrast, sleep deprivation resulted in longer steps, increased reliance on the center of mass state in the anteroposterior plane, and decreased instantaneous gait stability. These results suggest that gait adaptations following sleep loss cannot be fully explained by reduced cognitive resource availability.

## Acknowledgments

This work was supported by the KU Leuven Special Fund (grant no. 3M220016 to T.d.B.). The authors would like to thank all contributing students for their help with data collection and Prof. Geneviève Albouy for the insightful discussions.

## Notes

### Competing Interest Statement

The authors have declared no competing interest.

