## Supplemental Table 1 for "Effects of one night of sleep deprivation on single- and dual-task gait"

**Table S1**. Cohen’s d of pairwise comparisons

|  | **SDEP vs CON** | | **Single vs dual-task** | |
| --- | --- | --- | --- | --- |
| Outcome | **ST** | **DT** | **CON** | **SDEP** |
| Step time (s) | -1.354 | -1.783 | 0.773 | 0.344 |
| Step time variability (s) | -0.175 | -0.251 | 1.337 | 1.262 |
| Step length (m) | -1.190 | -1.612 | 0.844 | 0.422 |
| Step length variability (m) | 0.250 | 0.487 | 0.453 | 0.690 |
| Step width (m) | 0.547 | 0.564 | 0.169 | 0.186 |
| Step width variability (m) | -0.055 | -0.697 | 0.788 | 0.146 |
| R^2^_ML_ (%) | -0.212 | -0.667 | -0.408 | -0.863 |
| FPE_ML_ (m) | 0.306 | -0.356 | 1.641 | 0.979 |
| R^2^_AP_ (%) | -0.778 | -0.334 | -0.130 | 0.314 |
| FPE_AP_ (m) | 1.167 | 0.115 | 2.789 | 1.737 |
| MoS_ML_ (m) | 0.874 | 0.659 | 0.078 | -0.137 |
| MoS_ML_ variability (m) | 0.608 | -0.069 | 1.492 | 0.815 |
